# Combined aptamer and transcriptome sequencing of single cells

**DOI:** 10.1101/228338

**Authors:** Cyrille L. Delley, Leqian Liu, Maen F. Sarhan, Adam R. Abate

**Affiliations:** Bioengineering and Therapeutic Sciences, University of California San Francisco, San Francisco, 94158 California, USA; California Institute for Quantitative Biosciences, University of California San Francisco, San Francisco, 94158 California, USA; Chan Zuckerberg Biohub, San Francisco, 94158 California, USA.

## Abstract

The transcriptome and proteome encode distinct information that is important for characterizing heterogeneous biological systems. We demonstrate a method to simultaneously characterize the transcriptomes and proteomes of single cells at high throughput using aptamer probes and droplet-based single cell sequencing. With our method, we differentiate distinct cell types based on aptamer surface binding and gene expression patterns. Aptamers provide advantages over antibodies for single cell protein characterization, including rapid, *in vitro*, and high-purity generation via SELEX, and the ability to amplify and detect them with PCR and sequencing.

## Introduction

Cellular differentiation restricts the genetic programs that cells may execute, endowing distinct functions and phenotypes [1, 2]. This enables important abilities, like the generation of tissues and organs; however, dysregulation of this system can lead to diseases, like cancer [3]. On the microscopic level, important biological structures comprise heterogeneous ensembles of cells working in coordination [4-7]. The immune system, for example, uses multiple cell types to elicit a response to exogenous threats, prevent autoimmunity, and establish long-term memory [8]. In cancer, heterogeneity occurs in advanced malignancies and imposes a significant barrier to cure by often ensuring that a fraction of cancer cells resist treatment [9-11]. For heterogeneous systems, mixed, multicell measurements do not allow resolution of different cells into their functional groups. High throughput single cell transcriptome sequencing [12-14] is an effective tool for deconvoluting heterogeneity because it provides ample information to identify cell type [15] and infer cell state and function [16]. Moreover, it leverages the capacity of modern sequencing to analyze tens-of-thousands of cells per experiment, allowing analysis of populations [17-19]. Nevertheless, gene expression is dynamic and can change due to biologically important or trivial events, making data interpretation challenging. Indeed, there is often poor correlation between transcript count and protein abundance, particularly when measured in single cells [20, 21].

As the extreme boundary of cells, the membrane carries a molecular fingerprint useful for identifying cell type and function. This information is often complementary to gene expression data, being encoded in the proteome and, thus, containing features not otherwise observable, like post-translational protein modifications. Consequently, for cell type discrimination, surface profiling with antibodies and fluorescence-activated cell sorting (FACS) is the gold standard to classify cells into their myriad types via well-characterized biomarkers [22, 23]. However, the approach requires that each antibody be labeled with a unique fluorophore, limiting multiplexing to tens of antibodies. By swapping fluorophores with mass tags and using a mass spectrometer for the readout, over a hundred antibodies can be used [24, 25], although this is still far short of the tens-of-thousands of genes, splice-forms, and post-translational modifications actively used by organisms and available for characterization by antibodies. Ab-seq replaces the mass tags with nucleic acid sequence tags, using droplet-based single cell sequencing for the readout [26]. Because a sequence tag is encoded by its full nucleobase set, an astronomical number of sequence combinations are available for unique antibody labeling, shattering the multiplexing barrier. Moreover, the microfluidic approaches used to sequence single cell mRNA can be applied to the tags, allowing simultaneous surface and transcriptome profiling of single cells at high throughput [27, 28]. While this provides exciting opportunities for characterizing cells with paired gene expression and protein data, it requires access to high-affinity antibodies. Effective antibodies are available for common targets, but uncommon ones require custom generation, an involved process necessitating antigen purification [29, 30]. Antigen purification can be costly and labor intensive, and may not be possible depending on the native structure of the antigen [31, 32]. Once obtained, the antibodies must be conjugated to sequence tags, an additional step that can impact affinity. To enable simple and effective single cell surface profiling, an optimal approach would obviate the need for custom antibody generation and tag conjugation.

Here, we present Apt-seq, an approach to simultaneously profile the surfaces and transcriptomes of single cells using aptamers and single cell sequencing. Like antibodies, aptamers are affinity probes capable of specific binding to target epitopes [33-36], and aptamer binding can be multiplexed. Unlike antibodies, aptamers are nucleic acids in which the nucleobase sequence provides an intrinsic tag that can be read out via DNA sequencing, obviating the need for additional tag conjugation. Moreover, specific, high-affinity aptamers can be readily and inexpensively obtained with *in vitro* systematic evolution of ligands by exponential enrichment (SELEX) [34, 36, 37]. SELEX can be applied directly to living cells, avoiding antigen purification, and shortening the process from months to weeks [38-40]. This simplifies affinity reagent generation and enables new surface characterization only accessible to aptamers [41]. We demonstrate Apt-seq by using it to discriminate between cells based on aptamer binding and differences in gene expression.

## Results

### Aptamers allow single cell surface profiling via droplet barcoding and sequencing

Aptamers are nucleic acids that adopt a three-dimensional fold and bind specifically to protein epitopes and small molecules [34, 36]. Like antibodies, they can be used in combination for multiplexed characterization [42], while being easily identified via nucleic acid sequencing. To allow simultaneous sequencing of cell mRNA and aptamers, we polyadenylate the aptamers to mimic the structure of mRNA; this allows both to be captured and sequenced using identical poly-thymine primers (Fig. 1a). To label the cells with aptamers, the mixed aptamer library is incubated with a cell suspension, and unbound aptamers washed away (Fig 1b). To barcode the cells, we employ Drop-seq, a high throughput microfluidic approach [13], although other barcoding methods can also be used [12, 43-45]. In Drop-seq, cells are isolated in droplets with barcoded beads and lysis buffer (Fig. 1c) [13]. Upon lysis, aptamers and mRNA hybridize to poly-thymine barcode sequences on the beads (Fig. 1d), followed by demulsification, washing, and nucleic acid amplification [12, 46-48]. Amplification conjugates a unique barcode sequence to all aptamers and transcripts of a single cell, allowing material for many cells to be pooled, sequenced, and computationally deconvoluted by barcode. This provides, for every cell, paired aptamer and transcript reads (Fig. 1e) that are separated *in silico* (Fig. 1f and 1g).

**Fig. 1:**
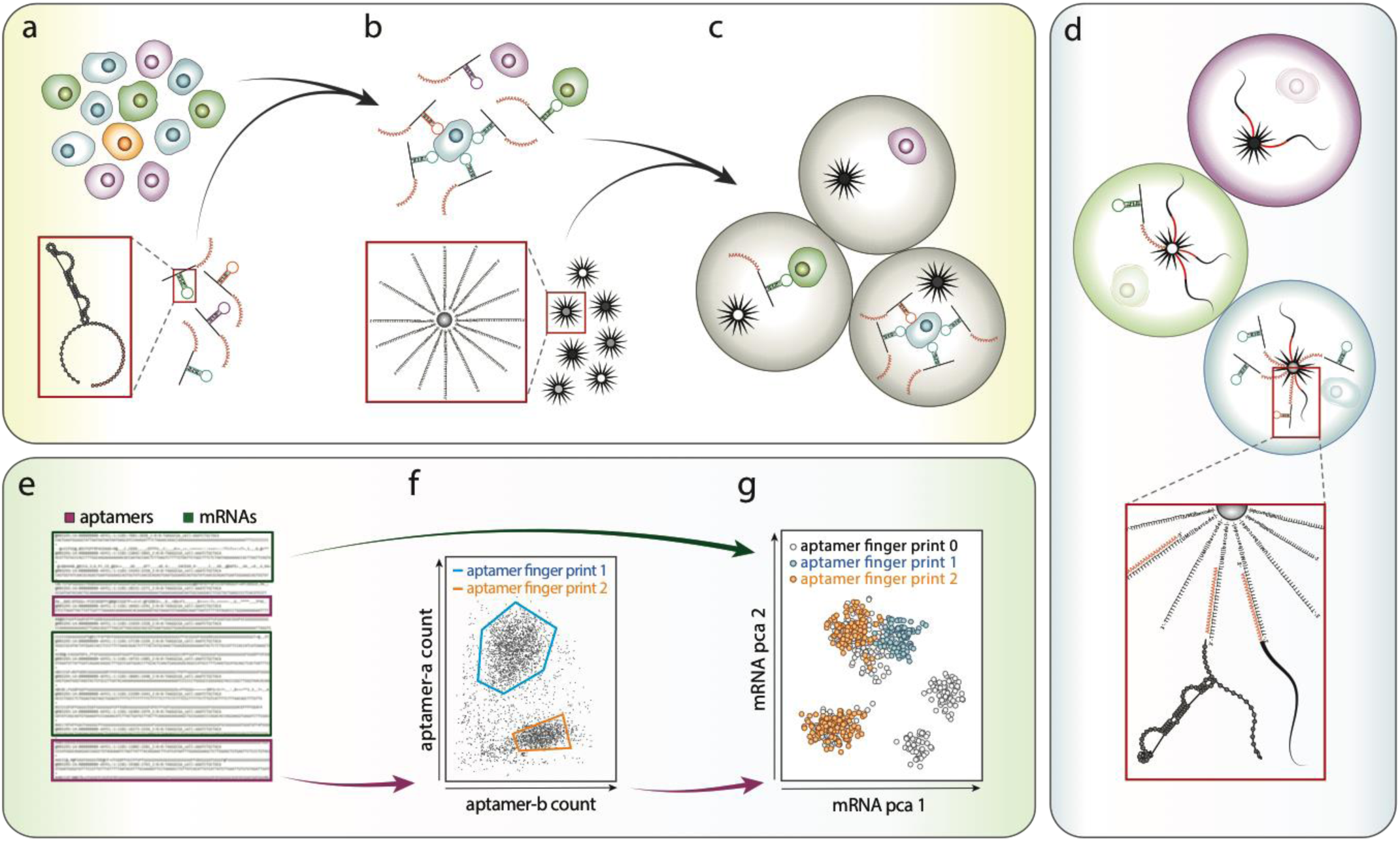
Principle of the Apt-seq workflow. **a** A heterogeneous cell sample is incubated with a diverse aptamer library containing a poly-A sequence on its 3’-end. **b** Cells expressing epitopes of interest are decorated by the corresponding aptamers in the library and non-binding aptamers are washed away. **c** Single cells of the washed cell suspension are co-encapsulated with beads carrying a unique DNA barcode in a microfluidic device. **d** Each droplet contains lysis solution to lyse cells. Aptamers and mRNA molecules can hybridize with the barcoding beads by means of their poly-A sequence. Using the barcode bead as a primer in reverse transcription and DNA polymerase reactions, the droplet-specific unique barcode is fused to the mRNA and aptamer, providing a cell specific identifier. **e** Pooling all beads after barcode fusion, sequencing their content in parallel, and deconvoluting aptamers and mRNAs, allows evaluation of epitope profiles in single cells **f. g** Since the cell-specific barcode is shared between aptamers and transcripts, the epitope data can be combined with the single cell transcriptome for further interdependent analysis.

### Polyadenylation does not impair aptamer function

For Apt-seq to be effective, the poly-adenylation required for paired transcriptome sequencing must not perturb aptamer binding [49]. To confirm this, we construct a library of five aptamers, TC01, TD05, TD08, TD09, and TE02, reported to bind Ramos cells with *K_d_* from 0.8 nM to 74.7 nM [50]. We also include TE17, sgc3b, and sgc8a aptamers that do not bind Ramos cells [42, 50, 51]. TD05, sgc3b, and sgc8a have reported protein targets, the membrane bound IgM, L-selectin, and PKT7, respectively [52-54]. To assess the impact of the poly-A tail on aptamer fold, we use RNAstructure [55], a secondary structure prediction algorithm, and predict the same fold for the aptamers with and without poly-A tail (Fig. 2a). To assess whether the tails interfere with binding, we synthesize all eight polyadenylated aptamers and apply them to Ramos and control 3T3 cells. The aptamers are incubated at equal molar concentration with either cell line, followed by five wash cycles and concentration estimation in the final wash supernatant and final cell suspension by qPCR. In agreement with previous studies, TD05, TD08, and TE02 are highly enriched in Ramos cell suspensions, while TD09 is moderately enriched. In contrast, TC01, TE17, and sgc8a are not enriched in Ramos cells, as expected (Fig. 2b) (Supplementary Fig. 1). Sgc3b remains below the detection limit for either cell line. Notably, although sgc8a is a reported binder of human T-cells [42], it enriches in mouse 3T3 cells. However, a previous study showed that both sgc8 and a PKT7 binding antibody can interact with other cell lines presumably devoid of PKT7 [56]. We conclude that poly-adenylating the aptamers does not affect fold or binding and that all except TC01 perform as previously reported in our hands.

**Fig 2:**
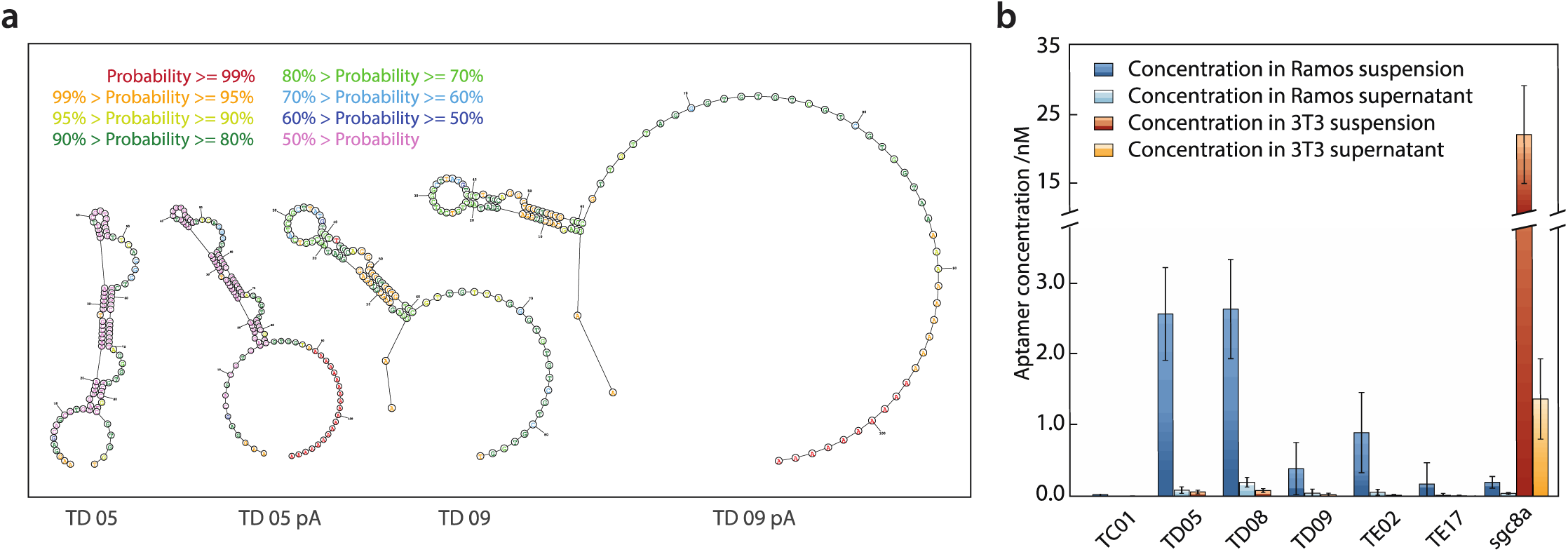
Influence of the 3’-poly-A tail on aptamer structure and function. **a** Predicted secondary structure of the aptamers TD05 and TD09 with and without poly-A tail. **b** The functionality of the aptamers TC01, TD05, TD08, TD09, TE02, TE17 and sgc8a, all modified with a 3’-poly-A tail is evaluated based on their ability to bind to Ramos or 3T3 cells. The concentration of the aptamers is estimated in the supernatant and cellular fraction after five wash cycles by qPCR. Each bar represents the mean of a technical triplicate and uncertainty is given as one sample standard deviation.

### Apt-seq provides independent information from RNAseq for inferring cell type

Inferring cell type from RNAseq data can be challenging due to the dynamic, complex, and multidimensional nature of gene expression. An advantage of Apt-seq is that it allows independent confirmation of cell type by aptamer binding. This can be used to support inferences from transcriptome data, or provide additional information for differentiating between related cells and states. To illustrate these benefits, we perform combined aptamer and transcriptome sequencing of suspended Ramos and 3T3 cells. After incubation and washing of the aptamers, we barcode the cells with Drop-seq. Drop-seq uses template switching to generate defined 3’-ends on cDNA, enabling subsequent PCR amplification. After amplification, cDNA is tagmented to generate ~500 bp fragments containing necessary sequencing handles. Aptamers lack the 5’-cap of mRNA and, thus, template switching is non-processive, preventing addition of the handles by this route. However, our aptamers are constructed with known flanking sequences, which we use for amplification and handle attachment. Additionally, the aptamers are short (below 200 bp), so that amplicons can be sequenced without tagmentation (Methods, Supplementary Table 1 and 2). These simplifications allow efficient, joint processing of aptamers and mRNA in the same barcoding reaction.

After barcoding, the nucleic acids of many cells are pooled and sequenced. We aim for about 200 cells and perform limited Illumina MiSeq sequencing collecting ~12 million paired-end reads. We computationally group reads to single cells using barcodes, and then to single molecules using unique molecular identifiers (UMIs). After processing, we obtain high-quality transcriptome and aptamer data for 58 cells, with >4500 unique reads per cell. More cells can be obtained by sequencing deeper and collecting more beads [13]. As expected, most reads belong to the transcriptome, and a smaller fraction to surface-bound aptamers (Fig. 3a). The aptamer profiles segregate into two groups, with TD08 and TE02, and to a lesser degree TD05 and TD09. The aptamer sgc8a is predominantly anticorrelated with this group. We obtain few reads for TC01, TE17, and sgc3b (Fig. 3b), consistent with our multicell qPCR results (Fig. 2b). Based on the qPCR results and because TD05 is a binder of the immunoglobulin heavy chain, we expect the first rectangular block to represent Ramos and the second 3T3 cells. Because these cells are from different species, cell type can be inferred by direct sequence analysis of cDNA. To verify our aptamer results, we thus evaluate the transcript reads of each cell. We order the transcriptome data using the same y-axis as the aptamers and again observe a clear block segregation, indicating that the gene expression of the two cell types is differentiated as expected. Based on transcript sequences, we confirm that the lower block corresponds to the human Ramos and the upper block to the mouse 3T3 cells (Fig. 3c).

**Fig 3:**
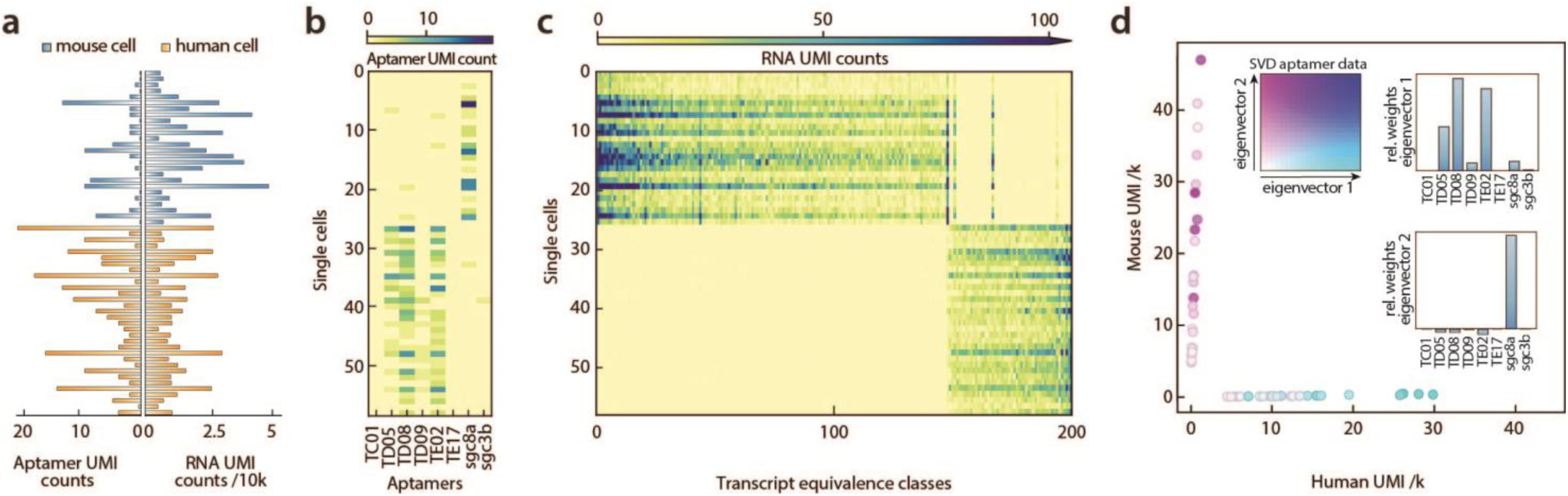
Mixed human-mouse two cell type experiment. **a** The number of unique aptamer reads per cell is drawn on the left. On the opposite side, all unique mRNA reads are displayed. The cells are ordered from bottom to top according to the number of assigned human mRNA reads divided by the total number of assigned mRNAs in decreasing order. **b** Two-dimensional histogram of counts per aptamer and cell. The order of the cells is the same as in **a**. **c** Two-dimensional histogram of the transcript counts for the 200 most variable transcript equivalence classes. The maximum read number is truncated at 100. The order of the cells is the same as in **a** and **b**. **d** Barnyard plot of the two cell transcriptome data. The first two principal components of the cell versus aptamer read matrix are superimposed in color space. Color code is given in the insert. The relative contribution of each aptamer to the first two principal components is given.

To confirm the relationship between expected aptamer and gene expression data, we construct a barnyard plot, displaying each cell as a 2D coordinate corresponding to the number of human and mouse transcripts identified within its barcode cluster (Fig. 3d). We extract the principal components of the aptamer read per cell matrix by calculating a singular value decomposition (SVD) (Fig. 3d, upper-left). This yields two eigenvector principle components, describing how the aptamers tend to correlate with each other. In agreement with the observed block structure of the aptamer profiles, we obtain an all positive first eigenvector with major contribution of the aptamers TD05, TD08, and TE02 (Fig. 3d, upper right) and a second eigenvector consisting primarily of sgc8a (Fig. 3d, lower-right). These eigenvectors are strongly anticorrelated, indicating that the aptamers cluster cleanly based on cell type. We use these eigenvectors to define a two-parameter coloring scheme, coloring each data point corresponding to a cell in the coordinate space according to its amplitudes along the two eigenvectors. Cells containing mostly human transcripts have amplitudes primarily along the first eigenvector (blue, Fig. 3c upper-right), while cells containing mostly mouse transcripts have amplitudes primarily along the second eigenvector (purple, Fig. 3d, lower-right). These results demonstrate that transcriptional signatures cluster with known patterns of aptamer binding and, more broadly, that aptamers can be used to label specific cell types in a format that can be read out with mRNA sequencing data.

## Discussion

While single cell mRNA profiling provides a valuable snapshot into the inner workings of cells, it is just one of many characterizations of the cellular space. Inaccessible from transcriptome data, the proteome contains traces of post-translational regulatory events and exhibits different temporal dynamics [57, 58]. Tapping into both domains, Apt-seq provides a strategy for highly multiplexed, single cell surface profiling simultaneous with the transcriptome. Because Apt-seq uses polyadenylated aptamer probes, it is compatible with most common single cell transcriptome sequencing methods, including Drop-seq, InDrops, and their commercial variants.

Apt-seq mirrors Ab-seq, and its successors CITE-seq and REAP-seq, in that it allows combined surface and transcriptome profiling of tens-of-thousands of single cells at high throughput [26-28]. However, aptamers provide practical and functional advantages over antibodies. In practical terms, aptamers are faster and cheaper to generate against many novel targets than antibodies. The *in vitro* SELEX process can be performed directly on living cells, obviating the need for antigen purification, which is not possible for many targets. Thus, Ab-seq may be preferred when the marker is best detected by a readily-available antibody, while Apt-seq may be superior for novel targets. Additionally, while antibodies require tag conjugation, aptamers can be used directly, since they are already sequenceable nucleic acids. In addition, patents on the original SELEX method have expired and new protocols have emerged that should expand the library of high-quality, specific aptamers [37, 59-61]. Moreover, aptamers can be synthesized with higher purity and reliability than antibodies through established chemical synthesis pipelines; this may allow aptamers to overcome the poor reproducibility that has plagued antibodies, and potentially achieve higher experimental reliability.

Apt-seq provides new capabilities for high throughput single cell characterization. In addition to scaling numbers of cells sequenced per sample, there is an increasing interest in scaling the number of samples sequenced in total. Ideally, many samples would be pooled and sequenced on a single run, but the need to index each one necessitates microfluidic barcoding of every sample, an expensive and labor-intensive process. Apt-seq can address this by labeling samples with unique surface aptamers that can be used independent of transcriptome data to batch samples together in one microfluidic barcoding run. Since after cell washing the free aptamer concentration is orders of magnitude below their dissociation constant, there should be minimal cross contamination between samples.

Aptamers and antibodies can bind internal epitopes of cells using fixation and permeabilization protocols; however, such binding may be better with the smaller aptamers, having a hydrodynamic diameter of ~2 nm compared to ~15 nm for antibodies [41, 62]. Indeed, many aptamers are readily internalized and staining of intracellular compartments can be more effective than with antibodies [63-65]. This will be important for combining intracellular proteomic and transcriptomic readouts [66]. Aptamers also enable measurement opportunities not possible with antibodies. For example, a raw aptamer library produced by cell-SELEX can be directly applied to cells, allowing different cell types to be inferred based on aptamer spectrum, rather than assembling an aptamer library from known binders. This should allow differentiation between cells even when distinct biomarkers are not known *a priori*. Additionally, multidentate, bispecific, or multipiece aptamers can be constructed by combining binding sequences in the same aptamer [67, 68], allowing assessment of epitope proximity in a way currently accomplished with fluorescence or proximity ligation assays, but in a format that can be highly multiplexed and performed simultaneously with cell surface and transcriptome profiling. Furthermore, riboswitch-like aptamers have a unique ability to bind small molecules and, thus, Apt-seq opens new possibilities for adding cellular metabolite characterization to simultaneous protein and nucleic acid measurements in single cells [69].

## Methods

### Aptamer structure prediction

The aptamer structure was predicted by the RNAstructure web-server (https://rna.urmc.rochester.edu/RNAstructureWeb/) with default parameters.

### Cell cultures

Ramos and 3T3 cells were cultured at 37°C in RPMI–1640 medium supplemented with antibiotics and 5% fetal bovine serum (FBS) in the presence of 5% CO_2_.

### Aptamer staining

Aptamers (from IDT) were folded as described before [50]. Briefly, they were diluted to 0.5 μM in aptamer folding buffer (F) (1x PBS, 5 mM MgCl_2_, 4.5 gl^-1^ glucose) and heated to 95°C for five minutes then cooled on ice for 10 min. The folded aptamers were pooled at a concentration of 62.5 nM each as stock solution.

3T3 and Ramos cells were resuspended in aptamer binding buffer (B) (buffer F supplemented with 1 mg/ml BSA and 0.1 mg/ml salmon sperm DNA). For the qPCR experiments either 3T3 or Ramos cells were diluted to about 6×10^5^ cells/ml and incubated with 31.25 nM of each aptamer. Cells were washed five times by centrifugation and resuspension cycles. For the single cell experiment 10^6^ Ramos cells where mixed with 3*10^5^ 3T3 cells and 31.25 nM of each aptamer in 1 ml buffer (B). The cell suspension was incubated for 30 min on ice and washed in ice cold buffer (B) for five centrifugation and resuspension cycles.

### Quantitative PCR

For each qPCR replicate either a suspension of about 4200 aptamer-stained cells or, in controls, the corresponding volume of supernatant was loaded. Aptamer specific primers (from IDT) were used to amplify individual sequences (Supplementary Table 2). Signal was detected on a QuantStudio5 (Applied Biosystems) qPCR machine with Syber green as reporter. A control experiment was performed to assess background priming on non-target aptamers and was found to be negligible (data not shown). Amplification standard curves were measured for all aptamer primer pairs in triplicate. The model *C*(*n*) = *C*_0_b*^n^*, where *C*(*n*) corresponds to the concentration after *n* PCR cycles and *C*_0_ to the start concentration, was log transformed and fit by linear regression to obtain the amplification rate *b* and the detection concentration *C*(*n_Ct_*), which is reached at the cycle threshold (*Ct*). Both calculated constants were used to estimate the aptamer concentration in the experiment. Uncertainty in the constants was propagated and final uncertainty of the concentration estimates is presented as one sample standard deviation with applied Bessel’s correction.

### Single cell experiment

The aptamer-stained cell suspension was diluted in ice cold PBS containing 0.1% BSA to a final concentration of 1.06*10^5^ cells/ml.

Single cell experiments were performed as described by [13] (online protocol: http://mccarrolllab.com/dropseq/), with the following modifications. About 2000 beads were used for further processing, corresponding to about 100-200 cells. After reverse transcription with Maxima H Minus Reverse Transcriptase (Thermo Fisher), which produces barcoded cDNA-mRNA hybrids and barcoded double-stranded aptamers, we amplified both libraries together by 13 cycles of PCR. DropSeq uses template switching to introduce the sequence ACTCTGCGTTGATACCACTGCTT at the 3’-end of the cDNA which allows use of a single primer (smart PCR) (Supplementary Table 2) to amplify the complete barcoded construct. Since template switching is much more efficient on 5'-capped mRNA, it only rarely happens on aptamers. Therefore, we use a specific reverse primer for each different aptamer flanking region (Supplementary Table 1), which we add to the PCR mix at a ratio of 1:20 compared to the smart PCR primer. Next DropSeq uses tagmentation (Nextera) to trim the transcripts to a mean length of about 500 bp and to introduce the sequence overhang GTCTCGTGGGCTCGG. Since the length of the aptamers is significantly shorter, tagmentation is not expected to act on them. To enable downstream joint processing of aptamers and transcripts we introduced the tagmentation overhang as part of the aforementioned specific primers. We processed the sample further as described [13]. The cDNA library was sequenced on a MiSeq device (Illumina) with a MiSeq Reagent Kit v3 (Illumina) in paired-end mode.

### Sequencing and data analysis

The python scripts for the evaluation of InDrops experiments [70] was adjusted to be applicable to the barcoding scheme of the DropSeq experiment. Without attempting any error correction, in a first pass all cellular barcodes with a count above 2000 where retained, which yielded 210 distinct barcodes. The corresponding sequences, containing transcripts and aptamers, were evaluated with Kallisto in the UMI-aware mode to generate a pseudo-alignment and to map transcripts to equivalence classes [71, 72]. To reduce the influence of stochastic fluctuations in the data, cells were only included for further analysis if they yielded a minimum of 4500 unique transcript reads; 58 cells passed this filter. For display purpose, the 200 equivalence classes with the highest variance were selected (Fig. 3c). A barnyard plot was generated based on equivalence classes that could be unambiguously assigned to either mouse or human only (Fig. 3d). Aptamer sequences were identified separately by the program cutadabt [73]. For each aptamer three overlapping 20mers were generated and each of these fragments was used as a cutadabt query and tested against all sequences. If a sequence contained a consecutive 20mer that matched the query with at most one base mismatch, the sequence was considered a read of that aptamer. The identified aptamer sequences were collapsed based on their UMI. SVD was calculated based on the mean-centered normalized cells versus aptamers count matrix. The color of the x-coordinate in the barnyard plot corresponds to the amplitude of the first eigenvector (cyan) and the y-coordinate to the inverse amplitude of the second eigenvector (magenta) (Fig. 3d).

## Data availability

The data that support the findings of this study are available from the corresponding author upon reasonable request.

## Acknowledgements

We thank John Haliburton for helping with Drop-seq setup and Anna Sellas of the Chan Zuckerberg Biohub for helping with the library sequencing. This work was supported by the Chan Zuckerberg Biohub, the National Science Foundation CAREER Award (Grant Number DBI-1253293); the National Institutes of Health (NIH) (Grant Numbers R01-EB019453-01, R01-HG008978 and DP2-AR068129-01); C.L.D. is supported by a Swiss National Science Foundation fellowship (Grant Number 175086).

## Contribution

A.R.A. conceived the project. C.L.D., L.L. and M.F.S. designed and carried out the experiments. C.L.D. evaluated the data. A.R.A. and C.L.D. wrote the manuscript.

